# IRAK1 is a critical mediator of low molecular weight hyaluronic acid-induced stemness in high-grade serous ovarian cancer

**DOI:** 10.1101/2023.09.25.559366

**Authors:** David Standing, Prasad Dandawate, Sumedha Gunewardena, Obdulia Covarrubias-Zambrano, Katherine F. Roby, Dineo Khabele, Andrea Jewell, Ossama Tawfik, Stefan H. Bossmann, Andrew K. Godwin, Scott J. Weir, Roy A. Jensen, Shrikant Anant

## Abstract

Advanced epithelial ovarian cancer (EOC) survival rates are dishearteningly low, with ∼25% surviving beyond 5 years. Evidence suggests that cancer stem cells (CSCs) contribute to acquired chemoresistance and tumor recurrence. Here, we show that IRAK1 is upregulated in EOC tissues, and enhanced expression correlates with poorer overall survival. IRAK1 and *BRCA1/2* mutation status are mutually exclusive. Moreover, low molecular weight hyaluronic acid (LMW HA), which is abundant in malignant ascites from patients with advanced EOC, induced IRAK1 phosphorylation leading to STAT3 activation and enhanced spheroid formation. Knockdown of *IRAK1* impaired tumor growth in peritoneal disease models, and impaired HA-induced spheroid growth and STAT3 phosphorylation. Finally, we determined that TCS2210, a known inducer of neuronal differentiation in mesenchymal stem cells, is a selective inhibitor of IRAK1. TCS2210 significantly inhibited EOC growth *in vitro* and *in vivo* both as monotherapy, and in combination with cisplatin. Collectively, these data demonstrate IRAK1 as a druggable target for EOC.

## Introduction

Recent advances in chemotherapy and surgical procedures have resulted in improvements in the overall survival of patients with epithelial ovarian cancer (EOC), yet it remains the deadliest gynecological malignancy with <25% surviving beyond 5-years ^1^. PARP inhibitors have demonstrated durable and highly effective antitumor responses in *BRCA*-mutated advanced EOC, which represents about 20% of the patients, though their effectiveness in *BRCA* wildtype and platinum-resistant EOC remain poor ^2^. The majority of EOC is high grade serous ovarian cancer (HGSOC) (1). The disease, which is thought to originate from the fimbriated end of the fallopian tubes, is frequently diagnosed at an advanced stage (stage III and stage IV), where the cancer has metastasized within the peritoneal cavity, leading to significant clinical complications ^3^. Delayed diagnosis has resulted in poor 5-year relative survival rates of ∼35% and ∼24% for stage III and IV disease, respectively ^3^. The poor survival statistics associated with advanced disease indicate that current therapeutic strategies including cytoreductive surgery and combination chemotherapy are insufficient. Hence, there is a critical need to better understand the biology of EOC to improve response to chemotherapy.

An explanation for the aggressive phenotype of HGSOC could be described by a population of cells, termed cancer stem cells (CSCs), that harbor high resistance to therapy via multiple mechanisms, including enhanced DNA repair, and increased expression of efflux multidrug resistance (MDR) transporters ^4,5^. One theory about the origin of CSCs is that they have arisen from differentiated cells that have acquired stem like properties due to a variety of factors including tumor microenvironment selective pressures ^6^. Interestingly, along the HGSOC progression timeline, tumor cells migrate from the fallopian tubes and metastasize into the peritoneum through the passive shedding of cells that form CSC-enriched multicellular aggregates or spheroids. These cells are exposed to an array of factors within the microenvironment, resulting in the clonal enrichment of resistant and aggressive cell populations. Therefore, these CSCs represent an opportunity for targeted therapy to enhance conventional therapeutic strategies.

Inflammation plays an important role in cancer initiation, development, progression, metastasis, resistance to chemotherapy, and stemness. Inflammatory processes such as repeated ovulation, endometriosis and pelvic infections has been linked to ovarian carcinogenesis ^7^. Toll/interleukin-1 receptor (TIR) signaling is a critical innate immune signaling pathway that engages IL-1R-associated kinase (IRAK1) to activate numerous downstream factors, such as NFκB, p38, JNK, and STAT3. These factors in turn, have implications in tumorigenesis, chemoresistance and stemness ^8–11^. Multiple mediators can activate the TIR pathway, such as IL-1β and microbe-specific conserved motifs known as pathogen-associated molecular patterns (PAMPs) ^12^. Amongst the PAMPs, lipoteichoic acid (LTA) and lipopolysaccharide (LPS), gram-positive and gram-negative bacteria respectively, are potent activators of TLR2 and TLR4.

In the present study, we report that IRAK1 is highly expressed in EOC and is associated with poor overall survival. Moreover, the *IRAK1* locus is amplified in a subset of EOC patient samples and is mutually exclusive from *BRCA1/2* and DNA damage repair gene mutations, suggesting a unique opportunity for targeted therapy when PARP inhibitors are ineffective. We demonstrate for the first time that IRAK1 is activated by low molecular weight hyaluronic acid (LMW HA), resulting in increased EOC growth and stemness. This finding is due to the significantly higher abundance of HA in malignant ascites when compared to either LPS or IL-1β. Following activation by LMW HA, IRAK1 induces stemness by regulating the expression of p38 MAPK and STAT3 proteins. In an *in-silico* screen, we have identified a novel IRAK1 inhibitor, TCS2210, which suppresses growth of EOC cells *in vitro*. This compound synergizes with cisplatin, thus providing a plausible rationale for targeting IRAK1 as a therapeutic strategy for EOC.

## Results

### TIR signaling is activated in CSC enriched spheroids and cisplatin resistant EOC cell cultures

EOC is plagued by high mortality rates due to the development of chemoresistance and tumor recurrence, which has been attributed, in part, to CSCs. Therefore, we sought to identify pathways involved in both the regulation of chemoresistance and stemness. To this end, we utilized A2780 cells and their isogenic cisplatin resistant cell line C30 as an established model for overcoming multidrug resistance ^13,14^. First, we confirmed cisplatin resistance of the C30 cell line by hexosaminidase assay. While cisplatin induced a dose- and time-dependent decrease in cellular proliferation of A2780 cells, its effects on C30 cells were minimal (**Figure 1A**). The calculated 72h IC_50_ was ∼2 μM and ∼100 μM for A2780 and C30 cells, respectively. The ability to thrive independently of substrate anchorage is a distinct property of CSCs. Cells grown as 3-D spheroids have been shown to enrich for CSC properties through the upregulation of CSC markers, drug efflux ABC transporters, and pluripotency factors ^15^ Additionally, cells grown in 3-D exhibit increased resistance to chemotherapy and increased tumorigenicity *in vivo* ^16,17^. Hence, spheroid cultures are accepted as a surrogate for stemness. We sought to determine if cells that have acquired resistance to cisplatin also have increased stemness. A2780 and C30 cells were grown in ultralow attachment conditions to evaluate spheroid forming potential. Cisplatin-resistant C30s formed significantly (p<0.001) more spheroids (∼20 spheroids) compared to cisplatin sensitive A2780 cells (∼7 spheroids), suggesting that stemness is linked to resistance (**Figure 1B**). To determine the changes in gene expression that led to cisplatin resistance and increased stemness, we performed total RNA-sequencing on A2780 and C30 cells, grown as either 2-D monolayers or 3-D spheroids. Differentially expressed genes were visualized in a cluster heatmap and volcano plots (**Figure 1C-1E**). Ingenuity Pathway Analysis (IPA) identified 43 pathways activated in C30 cells compared to A2780 cells (**Figure 1F**), and 18 pathways in 3-D spheroids compared to 2-D monolayers (**Figure 1G**). Of the top activated pathways, 2 pathways were commonly activated, the LPS/IL-1β mediated inhibition of RXR Function and RhoGDI Signaling (**Figure 1H**). This discovery suggested that these pathways may have dual roles in supporting both cisplatin resistance and stemness phenotypes. We further mined published RNA-sequencing datasets using the NCBI GEO databases to validate our findings. We determined that TLR/IL1 signaling genes were significantly upregulated in carboplatin resistant patient tumor compared to carboplatin sensitive tumor tissues (**Supplementary Figure 1A**). Moreover, TLR/IL1 genes were significantly upregulated in 3-D cultures of patient derived fallopian tube secretory epithelial cells compared to matched 2-D cultures (**Supplementary Figure 1B**). Collectively, these data confirm our findings, and suggest a pivotal role for TIR signaling in chemoresistance and stemness. In this regard, we also assessed the expression of stem cell markers and ABC transporters in our RNA-seq datasets and across NCBI GEO databases. We observed increased expression of the pluripotency factor *KLF4*, CSC markers *DCLK1* and *ALDH1A1*, and ABC efflux transporters *ABCB1*, *ABCC2*, and *ABCC3* in cisplatin resistant C30 compared to A2780 cells (**Supplementary Figure 1C**). Similarly, A2780 cells grown in 3-D exhibited increased *DCLK1*, *KLF4*, *ABCB1*, *ALDH1A1*, *ABCC2*, and *ABCC3* compared to cells grown in 2-D (**Supplementary Figure 1D**). Mining of NCBI GEO databases demonstrated similar results. We determined that pluripotency factors *KLF4* and *MYC* were upregulated in carboplatin resistant ovarian tumor tissue compared to sensitive tumor tissues (**Supplementary Figure 1E**). Furthermore, carboplatin resistance was associated with increased expression of ABC transporters, *ABCC3*, *ABCC5*, *ABCG1*, and *ABCG2* (**Supplementary Figure 1F**). Similar results were attained for fallopian tube secretory epithelial cells grown in 3-D cultures compared to 2-D (**Supplementary Figure 1G and 1H**). Collectively, these findings confirm a relationship between platinum resistance and the stemness phenotype.

**Figure 1.**
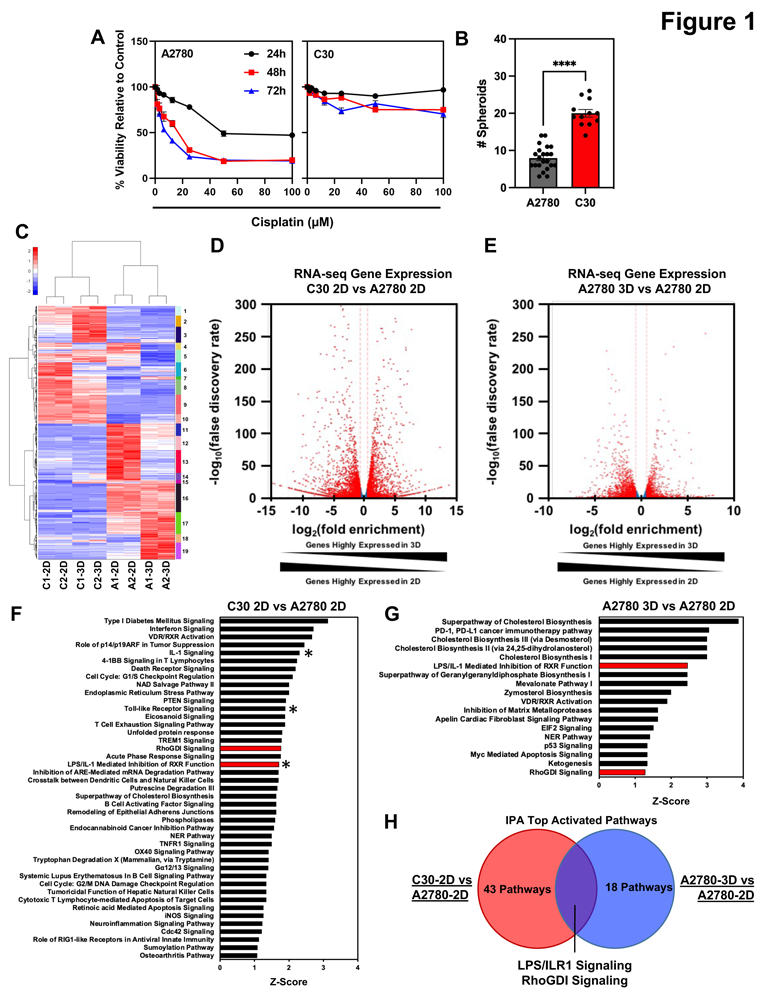
TIR signaling is activated in CSCs and cisplatin resistant EOC. (A) Viability assay of A2780 and C30 cells treated with cisplatin. (B) Quantification of spheroid formation for A2780 and C30 cells. (C) Hierarchal clustering heatmap of RNA sequencing data from A2780 and C30 cells grown as 2D and 3D cultures. (D) Volcano plot comparing log2 fold enrichment of genes highly expressed in C30 and A2780 cells grown in 2D. (E) Volcano plot comparing gene enrichment of highly expressed in A2780 cells grown in 3D and 2D. (F). List of IPA activated pathways in C30 cells. Red bars indicate commonly activated pathways. * Indicates TIR related signaling pathways. (G) List of IPA activated pathways in A2780 cells grown in 3D. Red bars indicate commonly activated pathways as in (F). (H) Venn diagram comparing IPA activated pathways from (F) and (G).

### IRAK1 is highly expressed in EOC

Given the activation of LPS/IL-1 signaling in both cisplatin resistant and 3D cultures, we probed the TCGA database to identify aberrantly expressed genes within the TIR signaling pathway in EOC data sets (**Figure 2A**). *IRAK1* was highly expressed across most samples analyzed in the Firehose Legacy dataset, comprised of 617 ovarian serous cystadenocarcinoma samples ^18^. We also determined copy number alterations (CNA) of TIR genes to understand mutation frequency in EOC. We observed that IRAK1 was altered in 51 samples (**Figure 2B**), which accounted for 9% of samples (**Figure 2B and 2C**). We also analyzed the copy number alterations of TIR genes across multiple TCGA datasets and determined that the majority of TIR genes were amplified in EOC (**Supplementary Figure 2A-C**). Overall, the number of CNA of TIR genes in EOC was observed in ∼25% of queried patients. In mining the Firehose Legacy serous ovarian adenocarcinoma dataset, we also identified that IRAK1 CNA were mutually exclusive from patients with *BRCA* and DNA damage-related repair gene mutations, defining a possible subset of EOC patients (∼25%) that may respond to TIR targeted therapies. These are the same tumors where PARP inhibitors would likely be ineffective (**Supplementary Figure 2D**). We proceeded to download the GDC TCGA OvCa dataset from UCSC Xenabrowser to further determine the impact of IRAK1 in EOC. The data was curated to remove redundant samples. Expression of *IRAK1* was then determined in normal tissue (Ctrl), primary tumor (PT), and recurrent tumor (RT) samples. *IRAK1* mRNA was significantly elevated in PT and RT compared to Ctrl (**Figure 2E**). In mining the NCBI GEO databases, we also determined that *IRAK1* mRNA expression was significantly upregulated in EOC compared to normal tissue (**Supplementary Figure 2E**). Upregulation of *IRAK1* was also observed in carboplatin resistant tumor tissue compared to sensitive tissue of OvCa patients (**Supplementary Figure 2F**) and A2780 cells exhibiting platinum resistance (**Supplementary Figure 2G**). Next, we evaluated the association of *IRAK1* expression with clinical parameters, including overall survival (OS) and diagnosis age. High *IRAK1* mRNA expression was linked to poorer OS compared to low expressing samples (**Figure 2E**) and was further associated with a younger diagnosis age (**Figure 2F**). We next sought to validate IRAK1 protein expression in tumor tissues. For this, we accessed tissue microarrays (TMAs) available through the University of Kansas Biospecimen Repository Core Facility (BRCF) at KUMC. These TMAs contained tissue cores of matched primary tumor (primarily HGSOC), metastatic tumor, and non-neoplastic fallopian tube tissue with samples from 100 patients. Tumor tissue exhibited significantly higher IRAK1 protein expression compared to normal fallopian tube, as determined by a board-certified pathologist (**Figure 2G & 2H**). Collectively, these data corroborate our findings from RNA-sequencing and IPA analysis, demonstrating that IRAK1, a critical mediator of the TIR signaling pathway, is highly expressed and linked to poorer OS in EOC.

**Figure 2.**
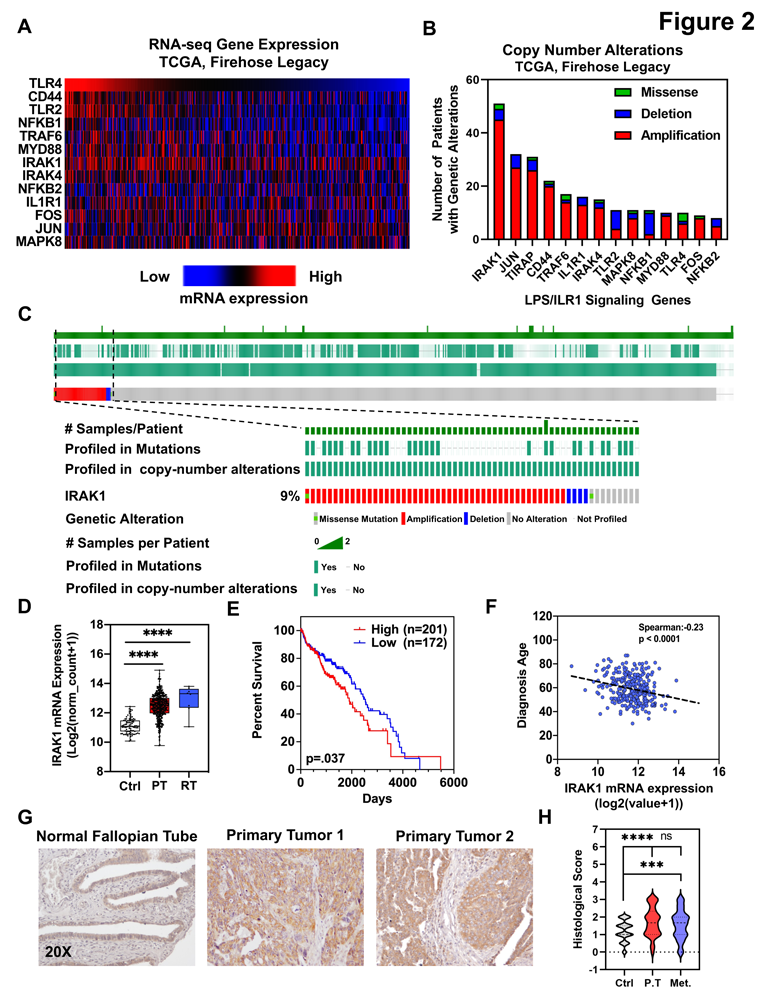
IRAK1 is upregulated in HGSOC. (A) Heatmap of mRNA expression of TIR signaling genes from CBioPortal TCGA, Firehose Legacy serous cystadenocarcinoma dataset. (B) Copy number alterations of TIR signaling genes from CBioPortal TCGA, Firehose Legacy serous cystadenocarcinoma dataset. (C) Oncoprint of TCGA, Firehose Legacy serous cystadenocarcinoma dataset. (D) Boxplot of IRAK1 mRNA expression in normal tissue (NT), primary tumor (PT) and recurrent tumor (RT) from TCGA Ovarian Cancer dataset. (E) Kaplan Meier survival curve of TCGA ovarian cancer dataset (High, n=201, Low, n=172, p=0.037). (F) Diagnosis Age versus IRAK1 mRNA expression of TCGA, Firehose Legacy ovarian serous cystadenocarcinoma dataset. Spearman correlation=-0.23, p<0.0001). (G) Representative immunohistochemistry for IRAK1 staining of normal fallopian tube and primary tumor from 2 patients from HGSOC TMA. (H) Histological staining score of IRAK1 in TMA, containing matched normal fallopian tube (ctrl), primary tumor (PT), and metastatic tumor (Met) from 100 patients with stage 3/4 HGSOC. ns=not significant, *** p<0.001, **** p<0.0001.

### Low molecular weight hyaluronic acid is abundant in malignant ascites and activates IRAK1 signaling in EOC cells

To begin evaluating the potential functional role of IRAK1 in the pathogenesis of ovarian cancer, we first performed western blot analysis to determine IRAK1 expression. IRAK1 was highly expressed in all EOC lines tested except PEO4 (**Figure 3A**). Interestingly, PEO4 cells, which were derived from a patient ascites at the time of relapse with cisplatin resistance, had the secondary *BRCA2* mutation, *i.e.,* 5193C>T (Y1655Y) that canceled the inherited mutation, *i.e.,* 5193C>G (Y1655X) and thus were BRCA2-proficient ^19^. A1847 and OVCAR8 were selected for further studies, as these are representative of the HGSOC histotype. IRAK1 was originally identified as a critical mediator of IL-1β induced activation of NF-κB through the IL-1R ^20^. Hence, we initially stimulated A1847 and OVCAR8 cells with IL-1β to validate functional IRAK1 signaling. We found that IL-1β increased IRAK1 phosphorylation at Thr209, a critical priming residue for IRAK1 activation (**Supplementary Figure 3A**). In addition, IL-1β induced phosphorylation of downstream canonical TIR signaling proteins, p38 MAPK and p65 (**Supplementary Figure 3A**). These data demonstrate that the pathway is active in these HGSOC models. We next sought to assess the role of IRAK1 activation in HGSOC from a biologically relevant perspective. Towards this end, we collected ascites fluid from 12 patients with advanced HGSOC and performed ELISAs for IL-1β and lipopolysaccharide (LPS), both potent activators of IRAK1 via IL-1R or TLR2/TLR4, respectively. Surprisingly, both IL-1β and LPS were expressed at low concentrations in malignant ascites (**Figure 3B**). Therefore, we investigated whether additional factors were present with the potential to activate IRAK1. Low molecular weight HA (LMW HA) has been previously shown to activate NF-κB through the binding of CD44/TLR2/TLR4 ^21^. Interestingly, CD44 is highly expressed in EOC primary tumors (PT), and recurrent tumors (RT) compared to normal tissue (Ctrl) based on GDC TCGA sequencing data (**Supplementary Figure 3B**). Furthermore, CD44 is highly correlated with IRAK1 mRNA (**Supplementary Figure 3C & 3D**), suggesting a novel link between CD44 and IRAK1 signaling. Hence, we performed an ELISA-like assay to determine if HA was present in malignant ascites and found that HA was present in high concentration relative to LPS and IL-1β, with a mean concentration of 110 ng/mL and max of ∼180 ng/mL (**Figure 3B**). This observation led us to investigate whether LMW HA could activate IRAK1. We stimulated A1847 and OVCAR8 cells with LMW HA (200 ng/mL). This rapidly induced phosphorylation of IRAK1 at Thr209 within 5 minutes of stimulation (**Figure 3C**). Next, we determined whether LMW HA promoted an important hallmark of cancer, namely stemness, since RNA-sequencing determined TIR signaling was activated in both cisplatin resistant and EOC CSC cell populations. For this, we stimulated A1847 and OVCAR8 cells with 200 ng/mL LMW HA and then evaluated spheroid forming potential. We observed that following incubation with LMW HA there was an increased number of spheroids with both cell lines compared to untreated controls (**Figure 3D & 3E**). To determine downstream signaling of LMW HA, we performed an unbiased phosphoproteomics analysis of cell extracts using a membrane-based antibody array containing 37 different kinases. We found that LMW HA treatment increased phosphorylation of canonical TIR factor p38-alpha, and non-canonical STAT3 and STAT5 (**Figure 3F**). We confirmed these findings by western blot analysis, in which LMW HA increased phosphorylation of STAT3, and p38 MAPK within 5 minutes (**Figure 3G**). We also observed increased expression of Myc, a STAT3 target gene (**Figure 3G**). Collectively, these data suggest a novel relationship between CD44, IRAK1, STAT3, and MYC in EOC.

**Figure 3.**
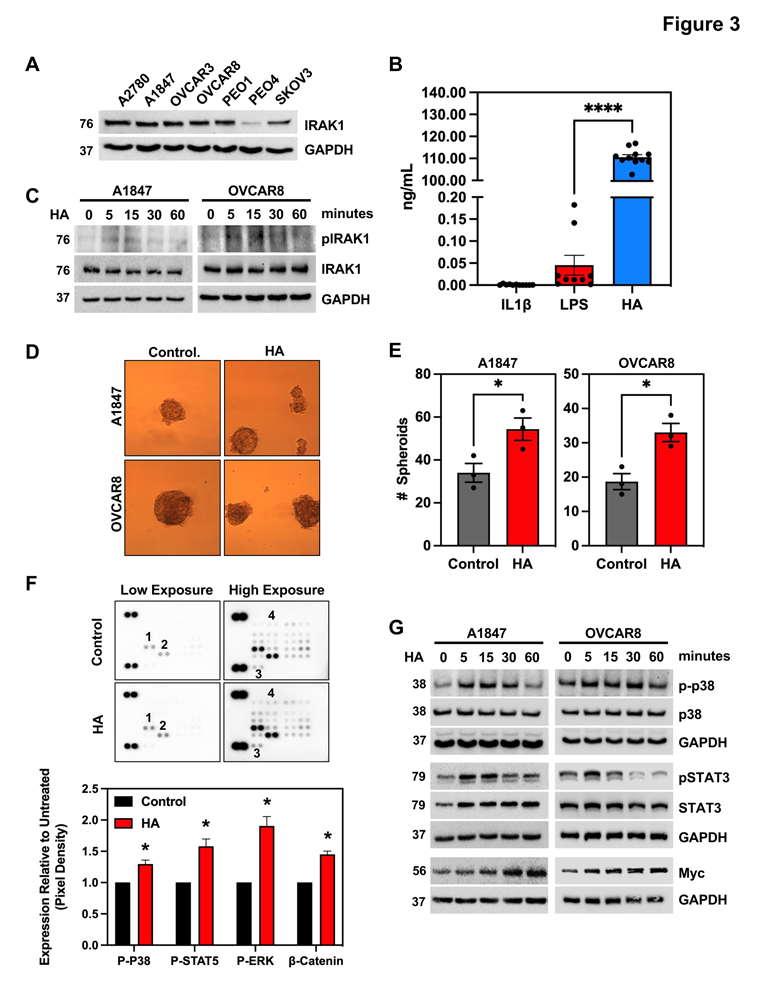
LMW HA activates non-canonical IRAK1 signaling and stemness. (A) Western blot for IRAK1 across EOC cell lines. (B) Quantification of ELISA for IL1β, LPS, and HA in malignant ascites from HGSOC patients (n=11). Data is represented as mean ± SEM (C) Western blot time course for pIRAK1 (T209) and total IRAK1 in A1847 and OVCAR8 cells following stimulation with LMW HA (200 ng/mL). GAPDH served as an internal control. (D) Representative images of spheroid formation assay in A1847 and OVCAR8 cells either unstimulated or stimulated with LMW HA for 14 days. (E) Quantification of (D). Data is represented as mean from 3 independent experiments ± SEM. (F) Representative image of low and high exposures of immunoassay-based kinase array following stimulation with or without LMW HA for 15 minutes. 1. pSTAT5; 2. pp38; 3. β-Catenin; 4. pERK. (G) Quantification of pixel density of (F) relative to untreated control. (H) Western blot of A1847 and OVCAR8 time course assessing phosphorylated and total expression of p38 and STAT3 following stimulation with or without LMW HA. The STAT3 target gene, *MYC*, was also assessed. * p<0.05, **** p<0.0001.

### IRAK1 is a critical factor that supports HGSOC tumor growth and stemness

To determine the role of IRAK1 in HGSOC, we transduced A1847 cells with either a non-targeting scrambled (Scr) construct or specific shRNAs targeting the *IRAK1* coding region and 3’UTR. *IRAK1* knockdown (KD) resulted in the downregulation of phosphorylated canonical proteins p38 and p65 (**Figure 4A**). Next, to delineate *IRAK1* regulated genes, we took an unbiased approach and performed total RNA-seq. Analysis of the data confirmed different gene expression profiles between *IRAK1* Scr and IRAK1 KD cells based on hierarchal clustering (**Figure 4B**). Specifically, *IRAK1* KD in A1847 cells decreased the expression of genes associated with stemness and multi-drug resistance (MDR), namely *CD44*, *STAT3*, *NOTCH1*, *NOTCH3*, and *ABCC1* (**Figure 4C & 4D**). Based on these findings, we next evaluated the impact of *IRAK1* KD on HGSOC cell growth and stemness by assessing colony formation (CF) and spheroid forming potential (SFP), respectively. *IRAK1* KD in A1847 cells significantly reduced colony size (p<0.0001) and number (p<0.01) (**Figure 4E & 4F**), and significantly (p<0.0001) impaired SFP (**Figure 4G & 4H**). Although not statistically significant, there was a trend towards smaller spheroids in the cells with IRAK1 KD. Collectively, these studies demonstrate that IRAK1 is critical for HGSOC proliferation and stemness. As further proof of principle, we next evaluated the effect of *IRAK1* KD on tumor growth *in vivo*. *IRAK1* Scr or *IRAK1* KD A1847 cells were injected intraperitoneally (IP) in NOD-*scid* IL2Rγ^null^ (NSG) mice to replicate advanced peritoneal disease of HGSOC. Mice injected with *IRAK1* Scr cells developed extensive peritoneal tumor growth and malignant ascites in 5/5 mice (**Figure 4I & 4J**). On the other hand, *IRAK1* KD failed to form tumors in 5/5 mice, which was further accompanied by a lack of development of malignant ascites (**Figure 4I & 4J**, p<0.001). Tumor weight was additionally measured following microdissection of tumors (**Figure 4K**). These data confirm that IRAK1 is a critical factor for HGSOC tumor growth. Since we previously determined that LMW HA induced phosphorylation of STAT3 and induced *MYC* gene expression (**Figure 3G**), we sought to confirm that these observations were dependent on IRAK1, and not a secondary off-target effect of shRNA transduction. To this point, we treated *IRAK1* Scr and *IRAK1* KD cells with LMW HA (200 ng/mL). HA induced phosphorylation of STAT3 within 5 minutes in *IRAK1* Scr control cells (**Figure 4L**). On the other hand, in *IRAK1* KD cells, HA induced phosphorylation of STAT3 was attenuated (**Figure 4L**). We also observed a reduction in total STAT3 in IRAK1 KD cells, confirming RNAseq analysis. This finding suggests that IRAK1 can regulate transcription of STAT3 in addition to its utilization as a substrate. To next verify that LMW HA-enhanced spheroid formation is dependent on IRAK1, we stimulated *IRAK1* Scr and *IRAK1* KD cells in ultralow attachment conditions to assess SFP. LMW

**Figure 4.**
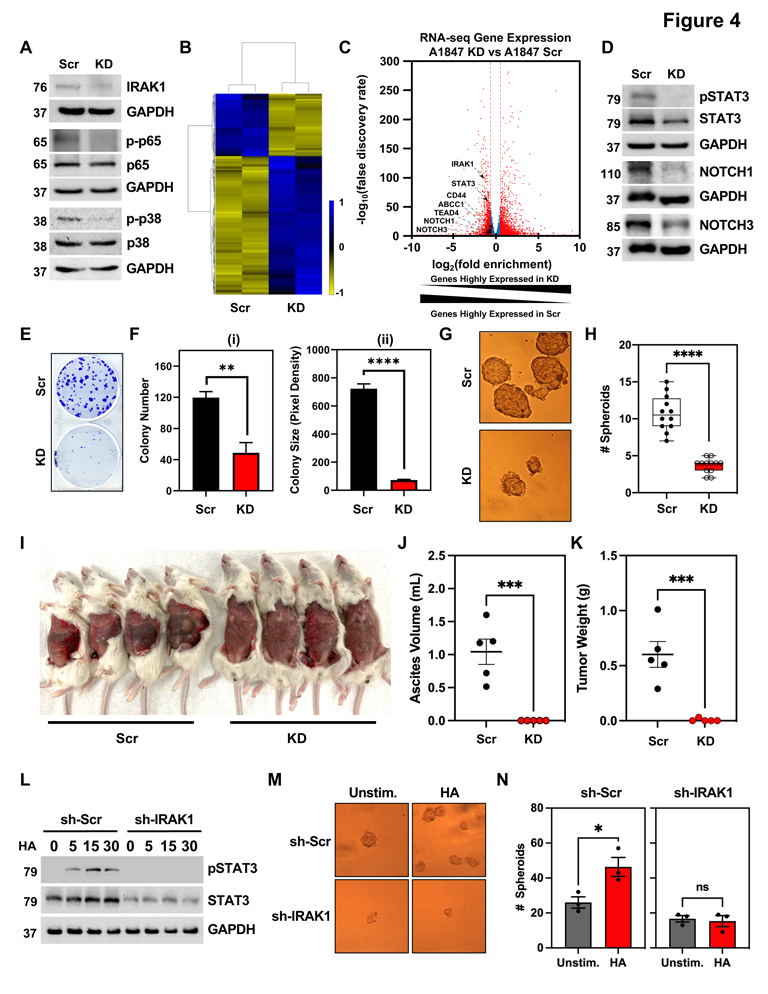
IRAK1 is critical for HGSOC growth and LMW HA induced stemness. (A) Western blot of IRAK1 in *IRAK1* knock down (KD) and *IRAK1* Scrambled (Scr) cells and phosphorylated and total p65 and p38. (B) Heatmap of RNA-sequencing comparing *IRAK1* KD and Scr gene expression. (C) Volcano plot of RNA-seq data from (B). Genes associated with stemness are labeled. (D) Western blot KD and Scr for pSTAT3, total STAT3, and cleaved Notch1 and Notch3. (E) Representative image of colony formation comparing KD and Scr. (F) Quantification of (E) for (i) colony number and (ii) colony size. Data are represented as mean from 3 independent experiments ± SEM. (G) Representative images of Scr and KD spheroid formation. (H) Quantification of spheroid formation from (G). Data are represented as box plots, min to max, all data points from 3 independent experiments. (I) Representative image of NSG mice injected IP with either IRAK1 Scr or KD cells. (J) Quantification of malignant ascites volume from IRAK1 Scr and KD groups (n=5 mice per group). Data represented as mean ± SEM showing all data points. (K) Quantification of tumor weight following microdissection from IRAK1 Scr and KD groups (n=5 mice per group). Data represented as mean ± SEM showing all data points. (L) Western blot time course of IRAK1 Scr and IRAK1 KD cells stimulated with LMW HA for phosphorylated and total STAT3. GAPDH served as internal control. (M) representative images of spheroid formation for IRAK1 Scr and KD cells stimulated with or without LMW HA for 14 days. (N) Quantification of spheroid number from (M). Data represented as mean ± SEM from 3 independent experiments. * p<0.05, ** p<0.01, *** p<0.001, ns=not significant.

HA significantly increased SFP in IRAK1 Scr cells (p<0.05); however, *IRAK1* KD impaired HA-induced SFP, confirming the critical role of IRAK1 in LMW HA-induced stemness (**Figure 4M & 4N**).

### TCS2210 is a selective inhibitor of IRAK1

Since IRAK1 KD impaired EOC growth, we next aimed to validate IRAK1 as a therapeutic target for EOC by characterizing the antitumor activity of a small molecule IRAK1 inhibitor. First, as proof of principle, we tested the Millipore Sigma IRAK1/4 inhibitor I (CAS-No 509093-47-4). This compound, however, had marginal anticancer activity in EOC cells. Hence, we sought to identify a selective IRAK1 inhibitor with high potency. We screened a library of mesenchymal stem cell modulating agents commercially available from Tocris using *in silico* docking (**Supplementary Figure 4A-D**). This method predicted potent interactions (BE < −9.0 kcal/mol) between IRAK1 and several compounds including CW008, TCS2210, Troglitazone, H89 and Kartogenin (**Supplementary Figure 4C & 4D**). While CW008 and H89 have been confirmed as PKA modulators ^22,23^, docking of CW008 predicted a single hydrogen bond interaction within the ligand binding domain, suggesting weak interaction. In addition, H89 has been shown to have pan-kinase inhibitory activity with preference for basophilic kinases ^24^. Troglitazone was removed from the US market in 2000, due to hepatoxicity ^25,26^. Kartogenin also displayed a single hydrogen bond with Gly-281, again suggesting a weak interaction. Therefore, these compounds were not considered for our studies. TCS2210 was predicted to bind within the “gatekeeper pocket” at Tyr-288, Leu-291, Asp-358, and Lys-239 with a binding energy of −10.4 kcal/mol (**Figure 5B**). To validate specificity and direct target engagement, TCS2210 was tested in the Eurofins KINOME scanMAX Active site Directed Competition Binding Platform, containing a set of 468 human kinases and disease relevant mutants. TCS2210 significantly mapped to a single target, IRAK1 (**Figure 5C**). To validate this, we performed cellular thermal shift assay (CETSA) and confirmed direct target engagement. We observed that TCS2210 protected IRAK1 from thermal denaturation, increasing the denaturation temperature from 46°C to 62°C (**Figure 5D**). We also performed surface plasmon resonance (SPR) studies, the gold standard for drug:target binding. SPR analysis revealed 1:1 binding kinetics with a K_d_ of 1.76 μM (**Figure 5E**). Collectively these data confirm TCS2210 binding to IRAK1. Based on these data, we sought to evaluate the ability of TCS2210 to block IRAK1 activation. We pretreated A2780 cells with 15 µM TCS2210 for 8 hours followed by stimulation with 200 ng/mL LMW HA. We observed that TCS2210 pretreatment blocked LMW HA induced phosphorylation of IRAK1 at T209, demonstrating efficacy as an IRAK1 inhibitor (**Figure 5F**).

**Figure 5.**
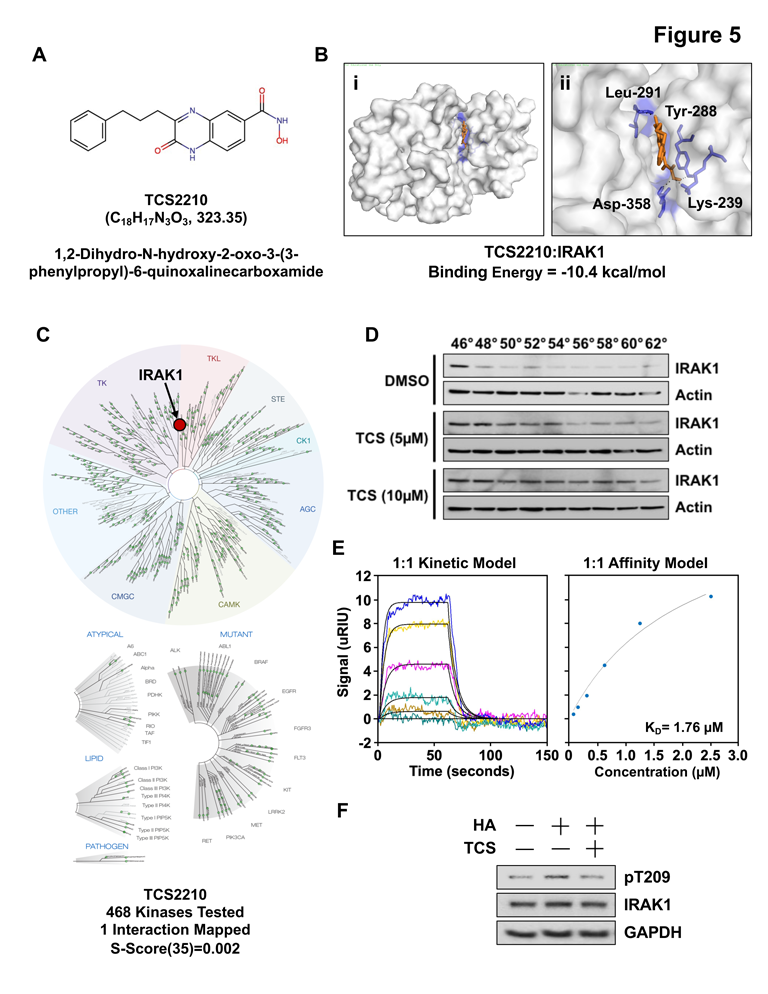
TCS2210 is a selective inhibitor of IRAK1. (A) Chemical structure of TCS2210 (B) *In silico* docking of TCS2210 with IRAK1: (i) space filled model (ii) zoomed in space filled model with predicted hydrogen bonding interactions. (C) Kinome tree interaction map of Eurofins ScanMAX analysis with TCS2210. (D) Western blot of CETSA assay for IRAK1 following incubation with TCS2210 or DMSO vehicle control. (E) SPR analysis of TCS2210 with active recombinant IRAK1 enzyme. Data are shown for 1:1 kinetic model and 1:1 affinity model. (F) Western blot of A2780 cells for pIRAK1 (T209) and total IRAK1 following preincubation with or without TCS2210, and stimulation with or without LMW HA.

### TCS2210 suppresses EOC growth and IRAK1 signaling

Having identified TCS2210 as a selective IRAK1 inhibitor, we next sought to confirm the targetability of IRAK1 for EOC. We treated A2780, A1847, OVCAR8, and OVCAR3 cells with increasing concentrations of TCS2210 for up to 72h. TCS2210 inhibited cell viability in all four cell lines in a dose- and time-dependent manner (**Figure 6A**). The calculated IC_50_ values for TCS2210 was ∼20 μM across all cell lines tested (**Figure 6A**). Furthermore, treatment of A1847 and OVCAR8 cells with TCS2210 significantly suppressed colony formation (**Figure 6B**). Even at ½ IC_50_, there was significant reduction in the number and size of colonies (**Figure 6C**). HGSOC cells treated with TCS2210 also exhibited increased AnnexinV/PI staining, revealing increased apoptosis (**Figure 6D**). We also observed that TCS2210 suppressed colony formation of A2780 and isogenic cisplatin resistant C30 cells (**Supplementary Figure 5A-C**). To evaluate the impact of TCS2210 on EOC stemness, we assessed SFP. TCS2210 significantly suppressed SFP in both A1847 and OVCAR8 cells at both ½ IC_50_ and IC_50_ doses (**Figure 6F**). Suppression was also observed in A2780 and C30 cells treated with TCS2210 (**Supplementary Figure 5C**). Functionally, we found that TCS2210 suppressed IRAK1 expression within 24h in A1847 and OVCAR8 cells (**Figure 6G**). In A1847 and OVCAR8 cells, we also observed decreased phosphorylation of STAT3 as well as reduction in the expression of the STAT3 target gene, MYC (**Figure 6G**). Collectively, these data confirm the results obtained with IRAK1 KD.

**Figure 6.**
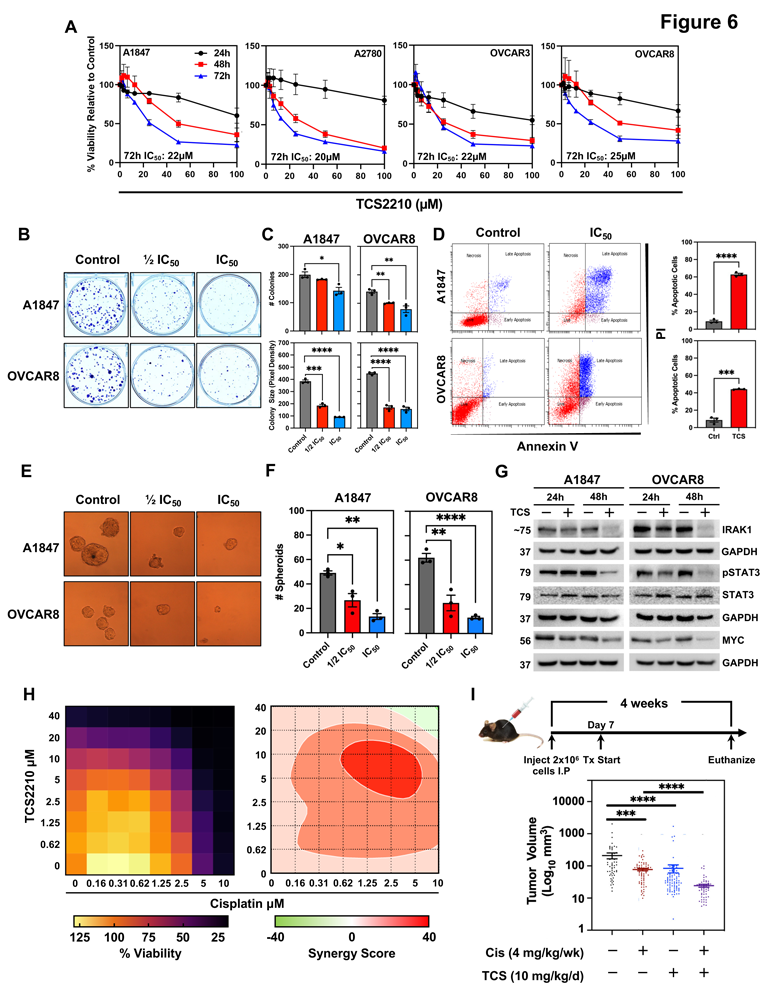
TCS2210 induces apoptosis of EOC cells. (A) Hexosaminidase viability assay in A1847, A2780, OVCAR3, and OVCAR8 EOC cell lines. Data are represented as means from 3 independent experiments ± SEM. (B) Representative images of colony formation in A18437 and OVCAR8 cells treated with TCS2210. (C) Quantification of colony formation number and size from (B). (D) Left: Representative image of scatterplot for Annexin V/PI apoptosis assay in A1847 and OVCAR8 cells treated with TCS2210. Right: Quantification of the percentage of apoptotic cells following treatment with TCS2210. Data are represented as mean from 3 independent experiments ± SEM. (E) Representative images of A1847 and OVCAR8 spheroid formation following treatment with TCS2210. (F) Quantification of spheroid formation from (E). Data are represented as mean from 3 independent experiments ± SEM. (G) Western blot of A1847 and OVCAR8 cells treated with TCS2210 for 24h and 48 hours. Immunoblots were performed for IRAK1, pSTAT3, STAT3, and MYC. GAPDH served as internal controls. (H) Left, heatmap of % viability for A1847 cells treated with various concentrations of TCS2210 and cisplatin. Right, synergy plot generated using SynergyFinder2.0 of A1847 cells treated in combination with TCS2210 and cisplatin. Data are represented as the mean from 3 independent experiments. (I) Top, experimental design of xenotransplant study. Bottom, quantification of tumor volume of micro-dissected tumors from mice (n=10 per group) injected IP with A2780 EOC cells and treated with either cisplatin (Cis), TCS2210 (TCS), or combination. Data represented as mean ± SEM showing all data points.

Since the majority of HGSOC patients show disease recurrence with presentation of chemoresistance to standard-of-care, we next sought to determine whether TCS2210 in combination with the standard-of-care agent, cisplatin would be a viable combination treatment strategy. In the *in vitro* proliferation assay, we observed that TCS2210 has strong synergy with cisplatin using the Zero Interaction Potency (ZIP) model by SynergyFinder (**Figure 6H**). The ZIP model overcomes many of the limitations of other synergy models; however, it implements the individual advantages of the Loewe and Bliss methods often used for this application ^27^. Subsequently, we determined the efficacy of TCS2210 as monotherapy and in combination with cisplatin in a peritoneal xenograft mouse model of advanced EOC (**Figure 6I**). Towards this, we first addressed the aqueous insolubility challenges with TCS2210 by preparing a β-cyclodextrin (CD) based formulation with a 1:1 molar ratio of TCS2210 and CD. To confirm that CD is non-toxic, we tested TCS2210, CD, and a 1:1 formulation (TCS+CD) by hexosaminidase assay in A2780 cells for 72h. We determined that CD had no inherent anti-cancer activity, while TCS2210 and TCS+CD inhibited A2780 cell growth (**Supplementary Figure 4G**). Moreover, TCS2210 and TCS+CD viability curves were nearly identical, indicating that CD had no activity altering effects. *In vivo*, co-administration of cisplatin and TCS2210 suppressed tumor volume of micro-dissected tumors by ∼50% (**Figure 6I**). The combination of cisplatin and TCS2210 achieved an even greater inhibitory effect, suppressing tumor volume by nearly 10-fold (p<0.0001). Collectively, these data demonstrate that IRAK1 is targetable by the small molecule inhibitor, TCS2210, resulting in EOC cells undergoing apoptotic cell death. Lastly, its synergistic activity with cisplatin indicates a plausible, promising new role for inhibiting IRAK1 for the treatment of EOC.

### TCS2210 nanoparticle suppresses EOC cell growth

Unfortunately, TCS2210 possesses poor physical and chemical properties required for development as a therapeutic agent, including poor aqueous solubility. To improve solubility, and thus, drug delivery, we developed a nanoparticle formulation of TCS2210. We generated a poly β-amino ester copolymer (PBAE) that was covalently linked to TCS2210, a CD44 targeting peptide, and a CD47 “don’t eat me” peptide, which we termed nano-TCS (**Supplementary Figure 5F**). Since CD44 is highly expressed in EOC, and is the cognate receptor for HA, we sought to enhance delivery of TCS2210 and selectively deliver this compound to ovarian cancer cells by incorporating a CD44 targeting peptide within the nanoparticle. Structurally, TCS2210 is linked to PBAE by ester bonds, which can be cleaved by intracellular esterases, improving compound stability and target delivery ^28^. We performed electron microscopy to evaluate nanoparticle size in an aqueous solution and determined that the average particle size was ∼35 nM (**Supplementary Figure 5G and 5H**). Lastly, we evaluated nano-TCS anti-cancer activity in A1847 cells by hexosaminidase assay. We determined that nano-TCS effectively suppressed viability of A1847 cells in a dose- and time-dependent manner (**Supplementary Figure 5I**), suggesting this drug delivery approach has merit in improving drug response.

## Discussion

HGSOCs are highly aggressive malignancies and the most common histotype of EOC with <25% surviving beyond 5-years ^3^. The disease is frequently diagnosed at an advanced stage (stage III and stage IV), where the cancer has metastasized within the peritoneal cavity, leading to significant clinical complications ^3^. Early in the EOC progression timeline, particularly as seen in HGSOC, cancer cells metastasize via a transcoelemic route within the peritoneum ^29^. Furthermore, these cells shed as 3-D spheroids enriched in CSCs, which are inherently more resistant to chemotherapy ^30^. Chemoresistance is commonly associated with advanced EOC cases, and this presents significant clinical challenges. Therefore, understanding the relationship between stemness and chemoresistance is necessary to improve treatment modalities and ultimately improve patient survivorship. We have demonstrated for the first time that IRAK1 is a critical mediator of stemness and EOC growth by regulating multiple pro-tumorigenic signaling pathways.

Current therapeutic paradigm involves cytoreductive surgery followed by the administration of systemic chemotherapeutic agents, including platinum and taxols. In a subset of patients with *BRCA1/2* mutations, resulting in HR deficiency, PARP inhibitors have shown durable and robust response; however, three-quarters of patients still develop resistance and hence, progress. Moreover, the use of PARP inhibitors for EOC patients who present with WT BRCA1/2, HR proficiency, and platinum resistant disease remains limited. We determined that IRAK1 amplification and expression is mutually exclusive from *BRCA1/2* mutation status, indicating that targeting IRAK1 may present a unique opportunity for therapy for patients that would otherwise have poor response to PARP inhibitors.

IRAK1 has been extensively characterized as a critical mediator of classical TIR signaling that activates NF-κB, and p38 MAPK controlled gene expression ^31,32^. In classical signaling, LPS or IL-1 bind to TLR4 or IL-1Rs and recruit Myddosome complex proteins MyD88, IRAK4, and IRAK1. IRAK1 is phosphorylated at Thr209 and Thr387 resulting in full activation of the kinase ^32^. We show for the first time that LMW HA activates IRAK1 in EOC. We have also determined that this in turn activates a non-canonical signaling axis centered around STAT3. A previous single study has shown that STAT3 is a substrate of IRAK1, albeit in murine splenocytes ^33^. The authors describe that LPS induced phosphorylation of STAT3 at both Y705 and S727, yet S727 was demonstrated as a direct IRAK1 phosphorylation site. In our own studies, we determined that STAT3 is phosphorylated at Y705 by LMW HA. Moreover, we show that this phosphorylation is dependent on IRAK1, since IRAK1 KD impaired STAT3 phosphorylation by LMW HA. While further studies are warranted on the mechanism of IRAK1 activation of STAT3, what is interesting is that this may be the strategic pathway for LMW HA induced stemness, since STAT3 is implicated in stemness and chemoresistance ^34,35^.

Our studies show that there are abundant levels of LMW HA in malignant ascites. On the other hand, factors that activate the canonical TIR signaling pathway, IL-1β and LPS are present at very low concentrations comparatively and may be insufficient to potently activate the pathway. This further demonstrates the clinical relevance of HA in EOC. In this regard, previous studies have also shown that the serum concentration of HA is significantly higher in ovarian cancer patients following carboplatin therapy. This was determined to be due to a resulting increase in the levels of the HA synthase (HAS2, HAS3), the enzyme that is essential for HA production ^36^. In the same study, the authors also demonstrate that HA treatment resulted in increased expression of drug efflux ABC transporters ABCB3, ABCC1, ABCC2, and ABCC3. Interestingly, this was only observed in cells expressing CD44, suggesting the signaling induced by HA following interaction with its cognate CD44 receptor ^36^. One of the earliest studies in ovarian cancer determined higher concentrations of HA at metastatic sites within the omentum ^37^. Furthermore, metastatic EOC cells shed as multicellular aggregates enriched in CD44+ cell populations ^38^. Collectively, this would imply that HA has implications in disease progression, contributing to the metastatic niche, and enhancing drug resistance at early stages of EOC development. Of note, the limitations of previous studies involved examination of HMW HA. HA is synthesized as large glycosaminoglycan polymers (up to 3400 kDa) in humans by HAS proteins ^39^. Typically, LMW HA is produced via catalysis of HMW HA by hyaluronidases (HYAL). Interestingly, HYAL 2 and HYAL 3 are overexpressed in OvCa, suggesting enhanced LMW HA production ^40^. Therefore, our studies examined a clinically relevant phenomenon, in which CD44 signaling is activated in response to LMW HA that is abundant within the tumor microenvironment of ovarian cancer patients.

Recently, there has been a renewed interest in TIR signaling for the development of novel therapies because inflammation plays important roles in tumorigenesis and progression of several malignancies. For instance, IRAK1 was characterized as a critical factor in metastasis and resistance to paclitaxel in triple negative breast cancer (TNBC) ^41^. Moreover, IRAK1 was associated with poorer OS. Both shRNA mediated knockdown and pharmacologic inhibition of *IRAK1* suppressed TNBC mammosphere formation. In a second study, use of the IRAK1/4 inhibitor (EMD Millipore CAS 509093-47-4) enhanced vinblastine activity against melanoma ^42^. In our studies, we also tested this reported IRAK1/4 inhibitor but determined marginal anti-cancer activity towards EOC cell lines. We initially rationalized that this was due to higher selectivity for IRAK4 based on compound specifications. However, since CD44 lacks a TIR intracellular domain, activation of IRAK1 would not occur through the IRAK4 containing myddosome complex. Rather, IRAK1 could be activated by PKC. It has been reported as a substrate for PKCδ ^43^, which is also activated by CD44 ^44^. Therefore, LMW HA induced phosphorylation of IRAK1 may be independent of IRAK4, especially in EOC. Most importantly, these studies indicate that there are IRAK4 independent mechanisms of IRAK1 activation, suggesting that compounds preferentially targeting IRAK4 may be inadequate. In this regard, a recent study also showed that knockout of MyD88 continued to demonstrate IRAK1 activation, suggesting an additional pathway to activate the protein ^45^.

Current crystal structures describe a critical structural element within IRAK1 called the regulatory spine ^46^. IRAK1 goes through a sequential phosphorylation event with the first phosphorylation at Thr209 for weak activation of the kinase, followed by Thr387 for full activation. The sequence surrounding T209 forms interactions with neighboring atoms, and a conformational change would be necessary for phosphorylation to occur. This conformational change within the protein is controlled by the regulatory spine, composed of residues F274, L263, F359, and H338 ^46^. Within the regulatory spine exists a structural element referred to as the “gatekeeper pocket”. The authors suggest that antagonist binding within the “gatekeeper pocket” inhibits the necessary conformational change for T209 phosphorylation. In our own modeling studies, we show an interaction between TCS2210 residues Y288, K239, D358, and L291 within the “gatekeeper pocket” of IRAK1. This suggests that TCS2210 likely inhibits the phosphorylation of T209 by obstructing conformational modifications within the regulatory spine of IRAK1. Certainly, additional biochemical studies are necessary to validate this hypothesis. Nevertheless, these findings provide a rational explanation for the mechanism of action for TCS2210 activity.

Lastly, we have developed a TCS2210 nanoparticle to enhance delivery of this compound and selectively target ovarian cancer cells. PBAE-based nanoparticles have been shown to be superior to stealth, pH-targeting, or antibody-based technologies, as these strategies have performed poorly in clinic ^47^. In our own studies, TCS2210 linked to PBAE can be cleaved by intracellular esterases, to protect the conjugated drug from premature enzymatic digestion and reduce off-target effects ^28,48^. We observed effective suppression of EOC cells *in vitro*. In the immediate future, we aim to validate the efficacy of nano-TCS2210 in suppressing IRAK1 activation and related signaling, as well as characterizing cellular uptake. In parallel with this approach to improve delivery, efforts are underway to generate and optimize IRAK1 inhibitors which possess properties necessary for drug development.

## Materials and Methods

### Cell Lines and Reagents

Cisplatin sensitive A2780 and their isogenic cisplatin resistant derivative, C30, as well additional human EOC cell lines A1847, PEO1, PEO4, OVCAR3, OVCAR8, and SKOV3 cell lines were generously provided by Dr. Andrew Godwin (University of Kansas Medical Center). All cell lines were grown in RPMI media (Corning, Tewksbury, MA) containing 10% FBS (Sigma-Aldrich, St. Louis, MO), 1% antibiotic/antimycotic solution (Corning, Tewksbury, MA) at 37°C with 5% CO_2_. Sections from paraffin embedded blocks containing matched adjacent normal, primary tumor, and metastatic tumor tissues of 100 de-identified EOC patient samples were obtained as a tumor microarray from the BRCF at KUMC. TCS2210 was purchased from Tocris (Minneapolis, MN). Hyaluronic acid (30,000-50,000 kDa) was purchased from Millipore Sigma (St. Louis, MO).

### Proliferation Assay

A2780, C30, A1847, OVCAR3, and OVCAR8 EOC cells (50,000 cells/mL) were plated in 96 well plates and allowed to grow for 24 hours in complete RPMI media. Cells were then treated with respective vehicle control (DMSO) or test compound (cisplatin, TCS2210). Cell viability was measured by hexosaminidase assay as previously described ^49^. Cell viability was calculated as a percent relative to untreated control.

### Spheroid Assay

Single cell suspensions of A2780, C30, A1847, and OVCAR8 cells (200 cells/well) were generated in 24-well ultra-low attachment plates (Corning, Lowell, MA). Spheroids were cultured in RPMI supplemented with EGF (20ng/mL), FGF (20 ng/mL), B27(10 mL), heparin salt (4 µg /ml) and pen/strep (1% v/v) (Invitrogen). Spheroids were allowed to grow for 5-10 days before being counted by a blinded observer. For HA related studies, spheroid media was diluted 1:5. Cells were then treated with a single dose of 150 µg of LMW HA. Spheroids were then allowed to grow for 14 days before being counted by an observer blinded to the treatment conditions.

### RNA-sequencing

The stranded total RNA-seq was performed using the Illumina NovaSeq 6000 Sequencing System at KUMC. Total RNA input ranging from 64.4 ng – 1000 ng was used to initiate the Illumina Stranded Total RNA Prep Ligation with Ribo-Zero Plus (Illumina 20040525) library preparation. Total RNA was processed using a probe based ribosomal reduction by Ribo-Zero Plus probes which were hybridized to rRNA targets with subsequent enzymatic digestion to remove the over abundant rRNA from the purified total RNA. The ribosomal depleted RNA fractions were sized by fragmentation, reverse transcribed into cDNA, end repaired and ligated with the appropriate indexed adaptors using the IDT for Illumina RNA UD unique dual indexes (UDI) (Illumina 20040553) to yield strand specific RNA-seq libraries. Following Agilent TapeStation D1000 ScreenTape QC validation of the library preparation and final library quantification by qPCR using the Roche Lightcycler96 with FastStart Essential DNA Green Master (Roche 06402712001), the RNA-Seq libraries were normalized to a concentration of 2nM and subsequently pooled for multiplexed sequencing on the NovaSeq 6000. The 100-cycle paired end sequencing run was performed using a NovaSeq 6000 S1 Reagent Kit v1.5 200 cycle (Illumina 20028318). Sequence data was converted to fastq files. The quality of reads was determined using FastQC software. Reads were mapped using the STAR software, version 2.6.1c ^50^. Transcript abundance estimates were calculated using the featureCounts software ^51^. Expression was normalized and differential gene expression calculations were performed using DESeq2 software^52^.

### Western Blot and Phospho-kinase Array

EOC cells (A2780, C30, OVCAR3, OVCAR8, PEO1, PEO4, SKOV3) were seeded in 10 cm tissue culture dishes at a density of 1×10^6^ cells. For time course studies, cells were plated in 100 mm tissue culture dishes in complete media. Media was replaced with serum free media for 24 hours before incubation with LMW HA (200 ng/mL). Cells were lysed, and protein isolated for downstream western blot analysis. Total protein lysate was subjected to polyacrylamide electrophoresis and transferred to PVDF membranes. Membranes were blocked in 5% milk and incubated with primary antibody (dilution 1:1000) overnight at 4°C. Protein expression was detected using HRP-conjugated secondary antibody (dilution 1:2000) and developed by ECL detection reagents (Amersham-Pharmacia, Piscataway, NJ). IRAK1 (4504S), pSTAT3 (9145S), STAT3 (9139S), MYC (9402S), KLF4 (12173S), p-p65 (13346S), p65 (8242S), p-p38 (9216S), p38 (8690S), Notch-1 (3608S), Notch-3(5276S) were purchased from Cell Signaling Technologies (Danvers, Massachusetts). IRAK1-T209 (SAB4504246-100UG) was purchased from Millipore Sigma (Burlington, Massachusetts). GAPDH (sc-32233) antibody was purchased from Santa Cruz Biotechnology (Dallas, Texas). Phospho-kinase array (ARY003C) was purchased from R&D Systems (Minneapolis, Minnesota). Lysates from A1847 cells treated with LMW HA (500 ng/mL) for 15 minutes were subjected to array analysis according to the manufacturer’s protocol.

### TCGA and NCBI GEO Database Analysis

Ovarian Serous Cystadenocarcinoma RNAseq data from The Cancer Genome Atlas (TCGA), Firehose Legacy, PanCancer Atlas, and Nature 2011 datasets, were downloaded using the cBioPortal for Cancer Genomics Browser (http://cbioportal.org). We also downloaded the TCGA GDC Ovarian Cancer dataset using the UCSC Xenabrowser (http://xena.ucsc.edu). We used the Firehose Legacy Dataset, to analyze the genetic alteration status and relative expression of critical genes involved in TIR signaling. PanCancer Atlas and Nature 2011 datasets were also interrogated to evaluate TIR signaling gene copy number alterations. We analyzed the mutation frequency of *IRAK1*, in comparison to mutation status and CNA of *BRCA1/2* and DNA damage repair genes (*RAD51, ATRX, SEM1, RPA1, NBN, ATR, ATM, CHEK1, CHEK2, FANCD2, FANCA, FANCC)* in the Firehose Dataset. We also evaluated diagnosis age relative to *IRAK1* mRNA. Using the GDC TCGA EOC database from Xenabrowser, we were able to analyze the relative expression of *IRAK1* mRNA across ovarian primary and recurrent tumors and normal samples. EOC datasets were curated to remove duplicate samples. For survivorship, a best expression cutoff was determined to stratify high and low IRAK1 expressing samples.

### Immunohistochemistry

Unstained tissue slide sections of five TMA blocks containing representative tissue cores (1 mm cores in triplicate) of matched benign fallopian tube, primary and metastatic tubo-ovarian high grade serous carcinoma tissues (n=100) were provided by the KUMC BRCF. Tissue microarrays were previously constructed by the BRCF staff using archival FFPE tissue blocks. The biospecimens and corresponding clinical information were de-identified to users. Paraffin-embedded tissues were de-paraffinized followed by antigen retrieval according to the manufacturer’s protocol. Briefly, tissue sections were incubated with UltraVision Hydrogen Peroxide block for 10 mins (Thermo Scientific). The slides were then incubated with IRAK1 primary antibody (Cell Signaling Technologies) overnight at 4°C. Slides were then washed and incubated with HRP Polymer Quanto for 10 mins before being developed with a DAB Quanto Chromogen-Substrate. Lastly, slides were counterstained with hematoxylin and eosin (H&E) before examination under a 20X objective. Slides were scored by a board-certified pathologist.

### ELISA for IL-1, LPS, HA

EILISA kits for IL-1, LPS, and HA were purchased from R&D Systems (Minneapolis, Minnesota). HA Ascites samples were obtained from the BRCF (University of Kansas Medical Center). and subjected to ELISA based quantification according to the manufacturer’s protocols. Briefly, the assays utilize a quantitative immunoassay technique. Standards, controls, and samples are added to wells. Unbound factors are washed away before an enzyme linked immunoglobulin is added. A substrate solution is subsequently added to the wells inducing a color change. A stop solution is then added, and the color is measured by a plate reader at the appropriate absorbance.

### IRAK1 Knockdown

shRNA targeting IRAK1 (NM_001569.3-2047s1c1, NM_001569.3-775s1c1, and NM_001569.3-2873s1c1) and a scrambled shRNA were purchased from Sigma Aldrich (St. Louis, Missouri). Lentiviral particles were made using the pLVX Advanced plasmid system (CloneTech Laboratories Inc, Mountain View, CA). Following transduction with lentivirus, A1847 cells expressing the transduced gene of interest were selected using 5 µg/mL Puromycin.

### Colony Formation

Cells were seeded at a density of 150 cells/mL in 6-well plates in complete growth media for 14 days. For TCS2210 treatment, colonies were treated 24 hours post-seeding for 72 hours. Media was replaced with complete growth media without drug. Media was subsequently changed on an as needed basis. After 14 days, cells were washed with PBS, and fixed with 10% neutral buffered formalin for 5 minutes. Colonies were then stained with 1% crystal violet (v/v) for 10 minutes. Colonies were then sufficiently washed before imaging and further analysis by ImageJ to quantify colony number and size.

### Animal Studies

For *IRAK1* KD xenografts, 2×10^6^ cells (A1847 Scr and *IRAK1* KD) were injected I.P in 5-week-old female NOD-scid IL2Rγ^null^ mice (n= 5 mice per group). Mice were evaluated for 28 days before euthanasia. Micro-dissected tumors were weighed. For TCS2210 and cisplatin anti-tumor efficacy studies, A2780 cells (2×10^6^ cells) were injected I.P in 5-week-old female nude mice. Tumors were allowed to form for 7 days before treatment was initiated. 10 mg/kg of TCS2210 was delivered as a β-cyclodextrin formulation (1:1 ratio) in PBS by I.P injection daily for 3 weeks. Additionally, cisplatin was delivered as a single dose weekly at 4 mg/kg. Combination treatment was also performed, using the dosing schedule described. Mice were euthanized on day 28 of study. Tumors were micro-dissected and weighed to quantify anti-tumor activity of treatment arms (n=10 mice per group). The Institutional Animal Care and Use Committee (IACUC) at the University of Kansas Medical Center approved all animal studies.

### Molecular Docking

AutoDock Vina software^53^ (Molecular Graphics Lab, Scripps Research Institute, http://vina.scripps.edu/) was used to analyze TCS2210 interactions with the 3D structure of IRAK1 (PDB ID: 6bfn). Molecular docking was performed using default parameters. Before docking interactions were calculated, total Kollman and Gasteiger charges were added to the protein and the ligand. The most stable compound:protein predicted conformation was selected for data presentation. This finding was based on the scoring function and the lowest binding energy ^54^.

### CETSA

TCS2210 interaction with IRAK1 was determined using the cellular thermal shift assay (CETSA) ^55^. Briefly, A2780 cells (10 × 10^6^) were treated in suspension with vehicle control (DMSO) or TCS2210 (5 and 10 μM) for 4 hours at 37°C and 5% CO_2_. The cells were then aliquoted by equal volume into PCR tubes. Following, cells were exposed to a temperature gradient (42-62°C) for 3 min. Cells were then lysed by repeated rapid freeze-thaw cycles. Total protein lysates were utilized in subsequent downstream immunoblot analyses to determine protein stabilization.

### Apoptosis Assay

To assess the induction of apoptosis by TCS2210, Annexin V/PI staining was performed. Briefly, 2 х 10^5^ cells (OVCAR8 and A1847) were seeded in 10 cm dishes for 24 hours. Cells were then treated with DMSO or TCS2210 at IC_50_ doses, as determined by hexosaminidase assay, for 72 hours. Cells were washed with PBS and stained as described in the manufacturer’s protocol with FITC conjugated Annexin V antibody and propidium iodide (PI). Cells were processed by flow cytometry to determine percent apoptotic cells.

### Combination Index (Synergy Studies)

A1847 cells were plated in 96 well plates at a density of 3000 cells/well. After 24 hours, cells were treated with increasing doses of cisplatin and TCS2210 for 72 hours. Plates were then developed by hexosaminidase assay, as previously described. Percent viability was calculated relative to untreated control. Data sets were then upload to the SynergyFinder2.0 (https://synergyfinder.fimm.fi/synergy/20221020000402075817/) server for analysis. Synergy was determined by ZIP method ^27^.

### Statistical Analysis

Data are reported as mean ± SEM. Data are representative of 3 individual experiments unless otherwise noted. Parametric, two-tailed t-test with Welch correction was performed to determine significance. For TCGA survivorship comparison between *IRAK1* high and low expressing groups, a log-rank (Mantel-Cox) test was used to assess differences. A best expression cut off was performed to stratify patient samples into high and low IRAK1 expressing groups. The FPKM value that yielded the maximal difference between survival at the smallest log-rank P-value was selected as the best expression cut off (0.037). All statistical analyses were calculated using GraphPad Prism software (version 9).

## Acknowledgements

This work was inspired by the late Dr. Shanthi Sitaraman, who died of HGSOC. If not for her encouragement and her commitment to overcoming ovarian cancer, this project could not have been possible. It is with great sadness and excitement that we dedicate this work to our dear friend. We would like to extend our gratitude to Dr. Michele Pritchard for her support and inspiration for pursuing HA in our studies. The authors have no competing interests to disclose. All data needed to evaluate the conclusions in the paper are present in the paper and/or the Supplementary Materials. RNA-sequencing data will be made available in the NCBI GEO Database. The research presented herein, received funding support from the University of Kansas Cancer Center Support Grant (P30 CA168524), Cancer Center Developmental Funds (to SA) and Biomedical Research Training Program (to DS).

Author contributions are as follows: DS curated data, performed data analysis, primary contributor to manuscript writing and figure preparation; PD curated docking data and analysis, prepared docking figures, manuscript textual editing/review; SG curated and analyzed RNA-sequencing data sets, manuscript textual editing/review; OC generated and curated data related to nanoparticle, manuscript textual editing/review; KR assisted with animal models, provided study oversight, manuscript textual editing/review; DK assisted with ovarian TMA acquisition, provided project oversight, manuscript textual editing; AJ provided project oversight, manuscript textual editing/review; OT analyzed IHC, manuscript textual editing/review; SHB provided nanoparticle development oversight, manuscript textual editing/review; AKG provided ovarian cancer cell lines, manuscript textual editing/review; SJW provided pharmacology oversight, manuscript textual editing/review; RAJ provided project oversight, manuscript textual editing/review, SA analyzed and reviewed curated data, assisted in manuscript writing and figure preparation, provided project oversight.

---

**Supplementary Figure 1.**
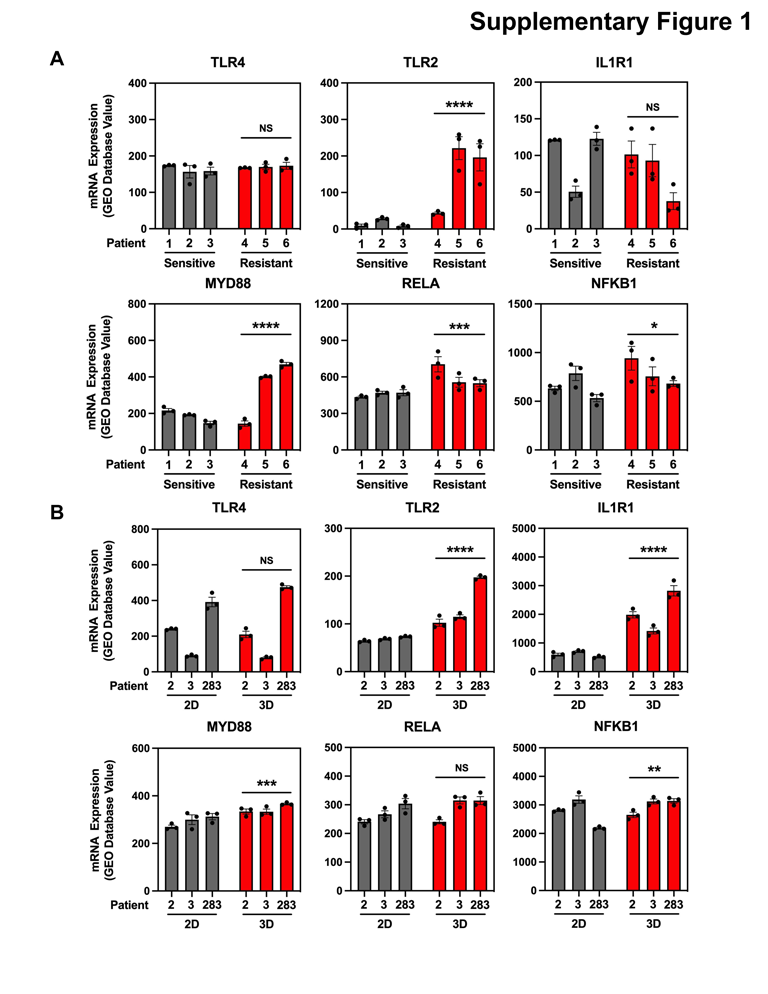
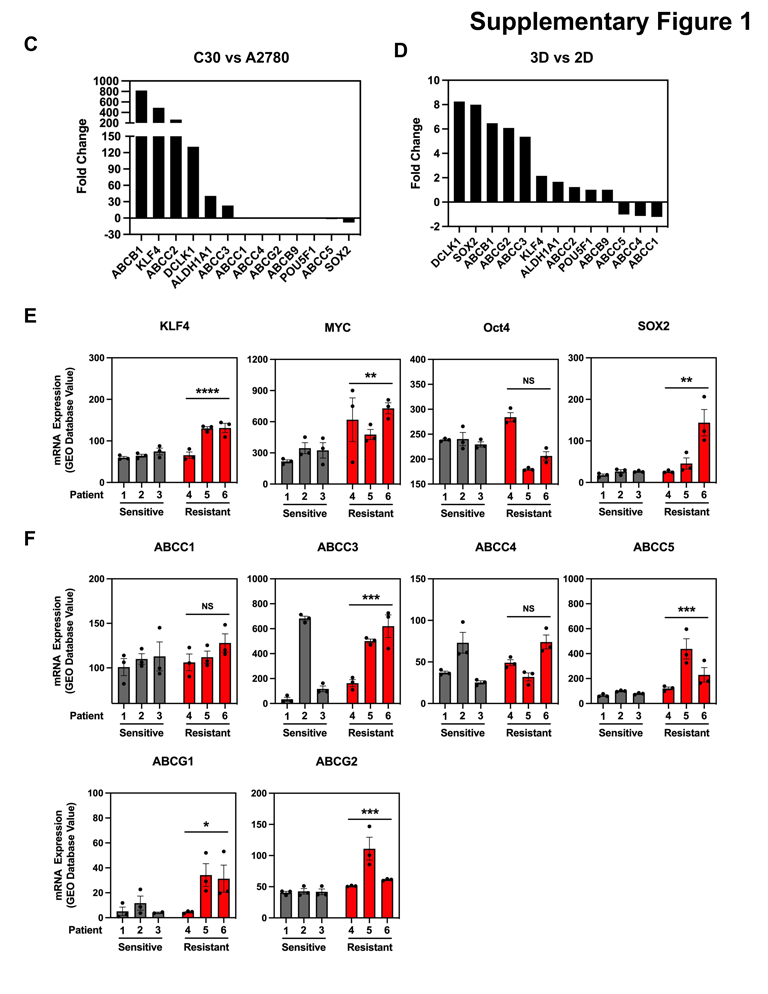
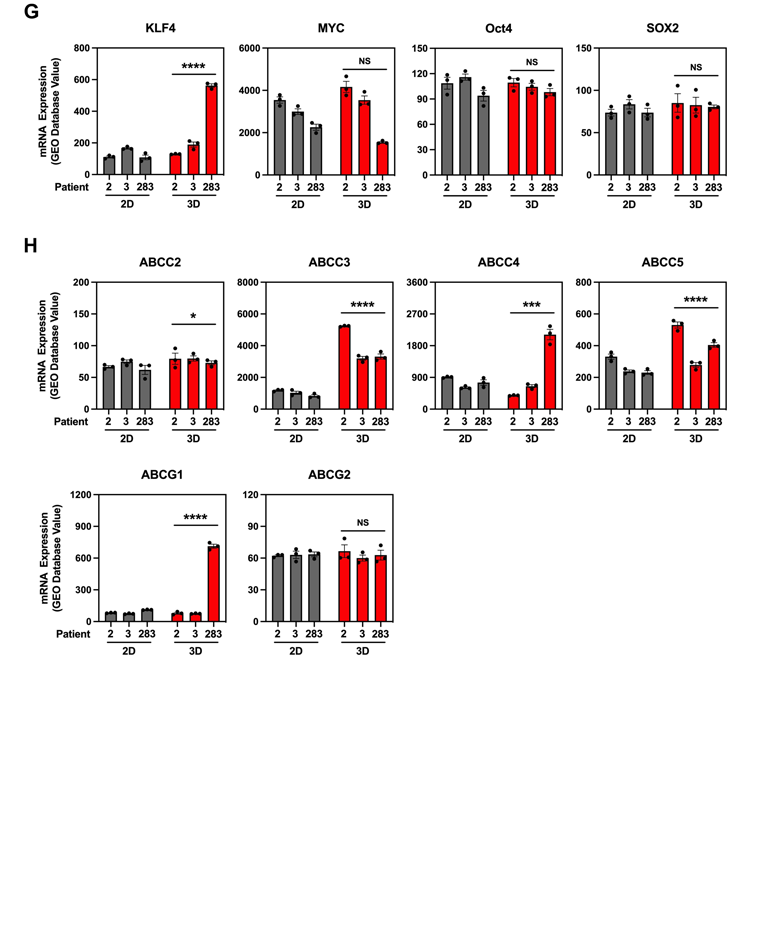
(A) Quantification of TIR signaling gene expression from NCBI GEO Database (GDS1381) comparing expression between carboplatin resistant and sensitive patients. (B) Quantification of TIR signaling gene expression from NCBI GEO Database (GDS5227) comparing 3-D culture effect on fallopian tube secretory epithelial cells. (C) Quantification of stemness and multi-drug resistant gene expression from RNA-sequencing in C30 cells compared to A2780 cells grown in 2-D cultures. (D) Quantification of stemness and multi-drug resistant gene expression from RNA-seq of A2780 cells grown in 3-D cultures compared to 2-D. (E) Quantification of IPSC stemness gene expression from NCBI GEO Database (GDS1381) comparing expression between carboplatin resistant and sensitive patients. (F) Quantification of multi-drug resistant gene expression from NCBI GEO Database (GDS1381). (G) Quantification of IPSC stemness gene expression from NCBI GEO Database (GDS5227) comparing 3-D culture effect on fallopian tube secretory epithelial cells. (H) Quantification of multi-drug resistant gene expression from NCBI GEO Database (GDS5227). Data are presented as mean showing individual technical replicates ± SEM from 3 independent patients. * p<0.05, ** p<0.01, *** p<0.001, **** P<0.0001, NS=not significant.

**Supplementary Figure 2.**
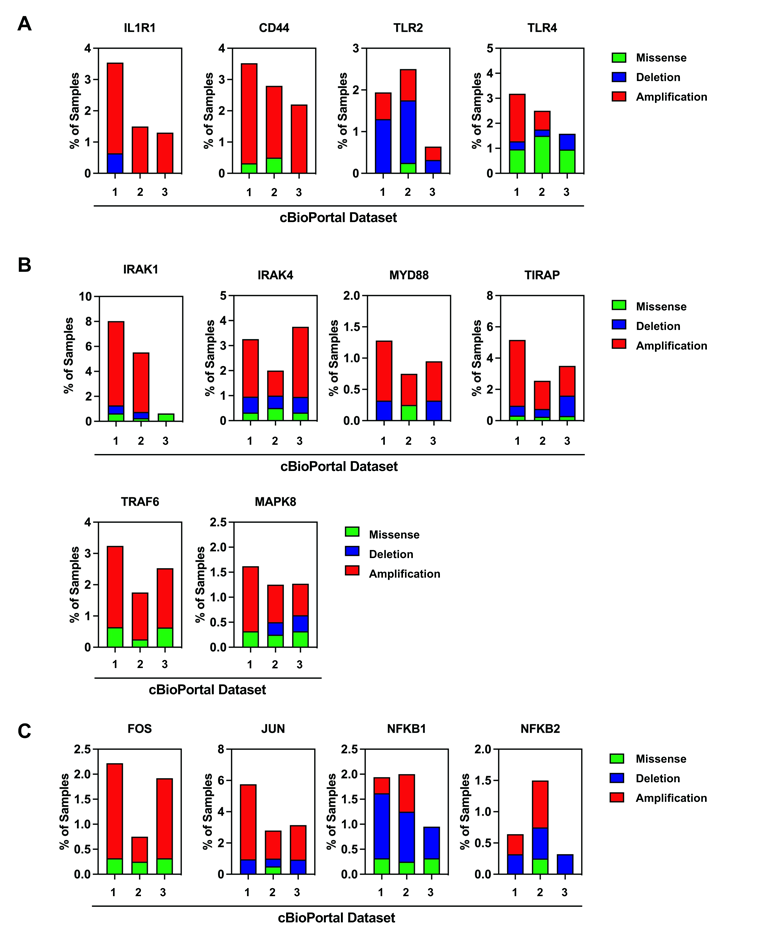
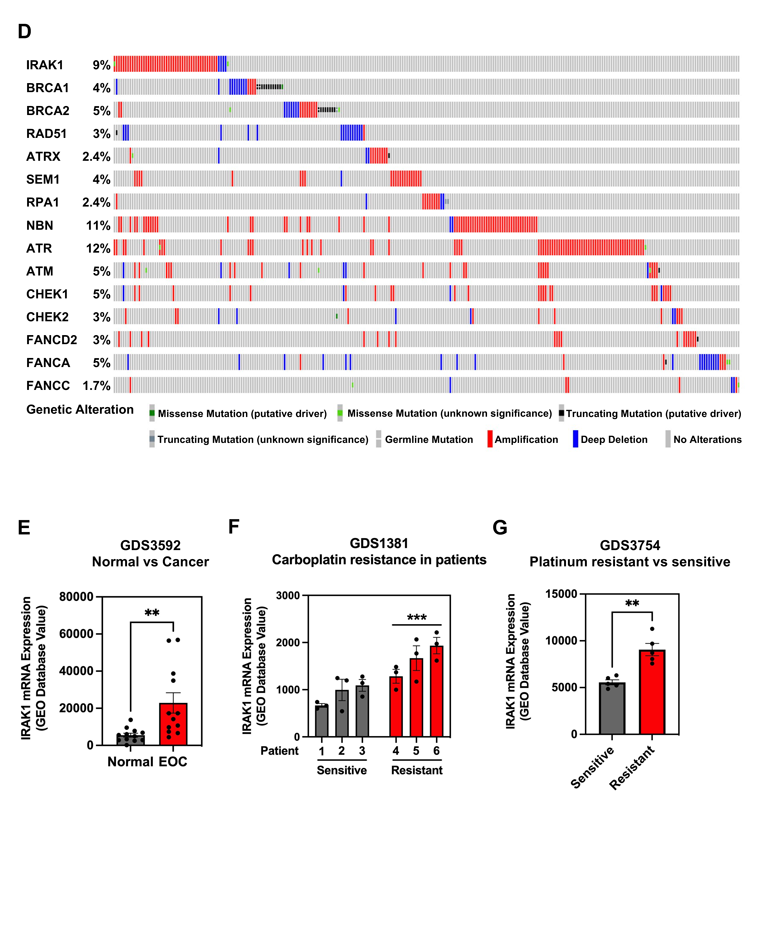
(A) Quantification of copy number alterations in TIR signaling receptors from 3 independent cBioPortal ovarian serous adenocarcinoma datasets. 1: Firehose Legacy 2: PanCancer Atlas 3: Nature 2011. (B) Quantification of copy number alterations in TIR signaling mediators as in (A). (C) Quantification of copy number alterations in TIR transcription factors as in (A). (D) Oncoprint of genetic alterations comparing IRAK1 and genes related to *BRCA* and DNA damage repair from cBioPortal ovarian serous cystadenocarcinoma Firehose Legacy dataset. (E) Quantification of IRAK1 mRNA expression from NCBI GEO Database (GDS3592) comparing ovarian normal surface epithelia and ovarian cancer epithelial cells. (F) Quantification of IRAK1 mRNA expression from NCBI GEO Database (GDS1381) comparing carboplatin resistance in patients. (G) Quantification of IRAK1 mRNA expression from NCBI GEO Database (GDS3754) comparing A2780 epithelial ovarian cancer cell line with acquired platinum resistance. Data are presented as mean ± SEM. Statistical significance was calculated by 2-way ANOVA ** p<0.01, *** p<0.001.

**Supplementary Figure 3.**
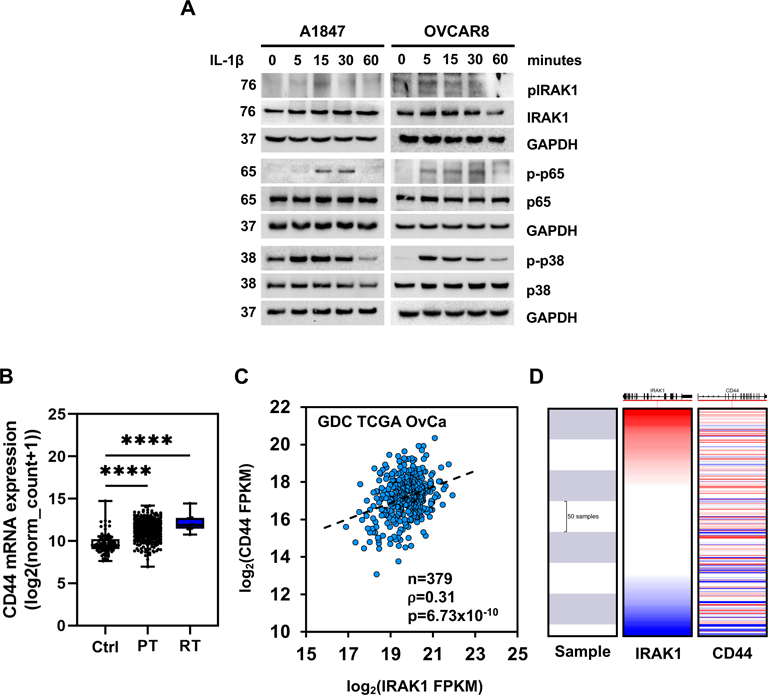
(A) Western blot time course in A1847 and OVCAR8 cells following stimulation with or without IL1β (10 ng/mL) for phosphorylated and total IRAK1, p65, and p38. (B) Quantification of CD44 mRNA from TCGA ovarian cancer dataset in normal tissue (NT), primary tumor (PT), and recurrent tumor (RT). Data is represented as a bloxplot, min to max, showing all data points. (C) Correlation of CD44 mRNA versus IRAK1 mRNA, represented as log2 FPKM from GDC TCGA EOC dataset (n=379). (D) Graphical correlation of (C). Red represents high expression. Blue represents low expression.

**Supplementary Figure 4.**
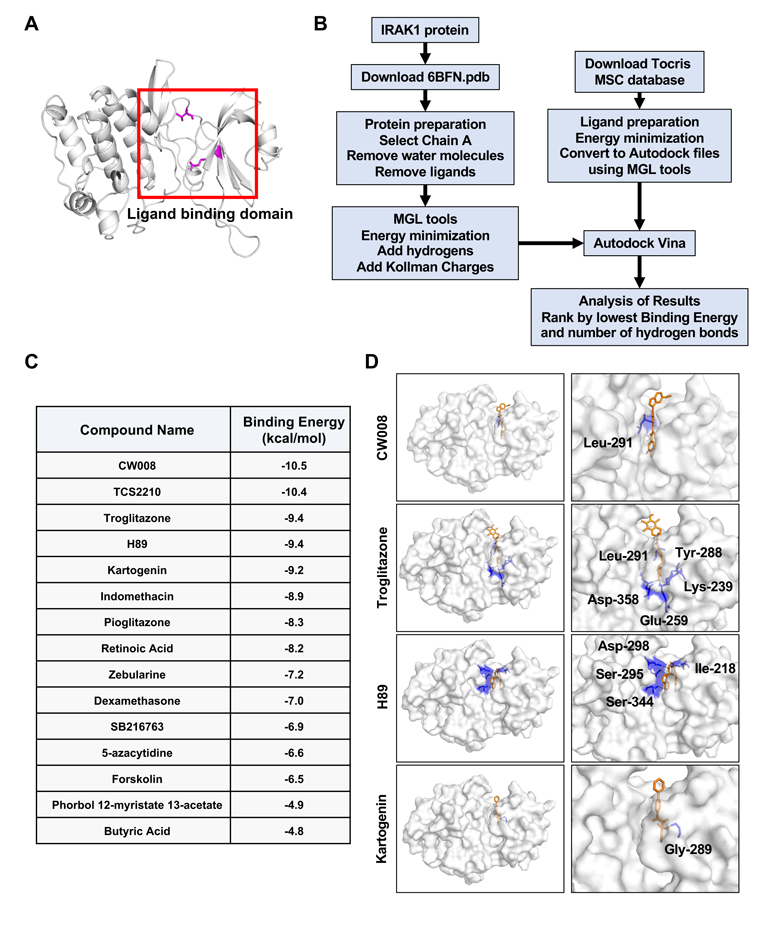
(A) Ribbon model of IRAK1 protein, highlighting domain used for *in silico* docking. (B) Schematic of docking workflow for *in silico* screening. (C) Summary of mesenchymal stem cell differentiating compound list from Tocris and binding energies following *in silico* docking screen with IRAK1 within the ligand binding domain. (D) Space-filled models of CW008, troglitazone, H89, and kartogenin binding within the IRAK1 ligand binding domain. Amino acids involved in hydrogen bonding are highlighted.

**Supplementary Figure 5.**
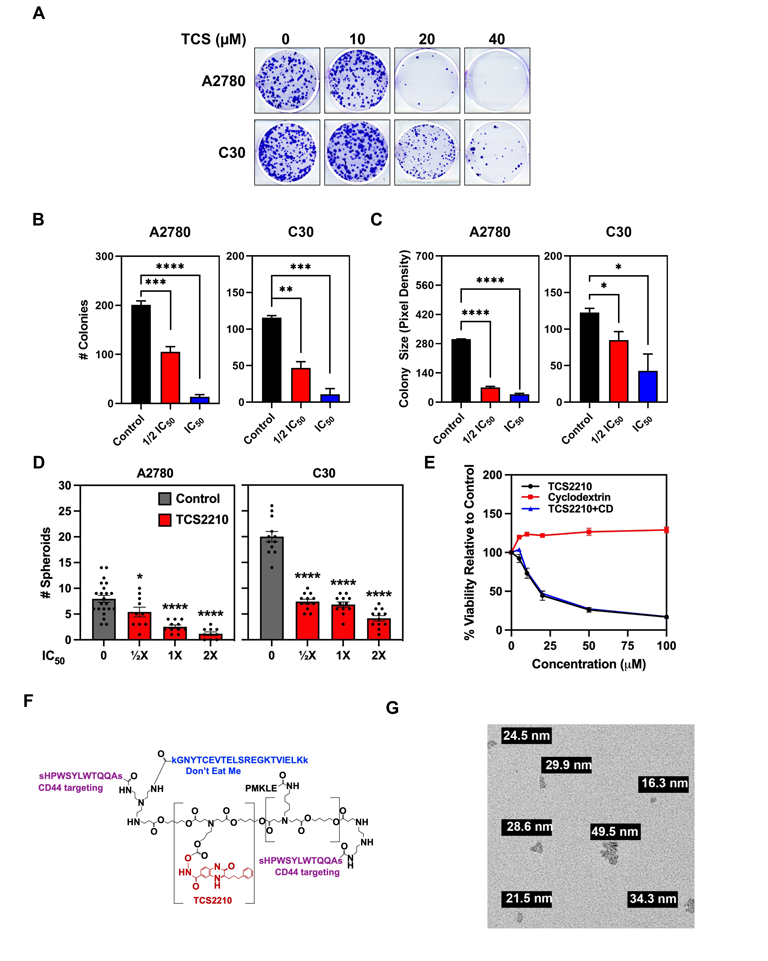
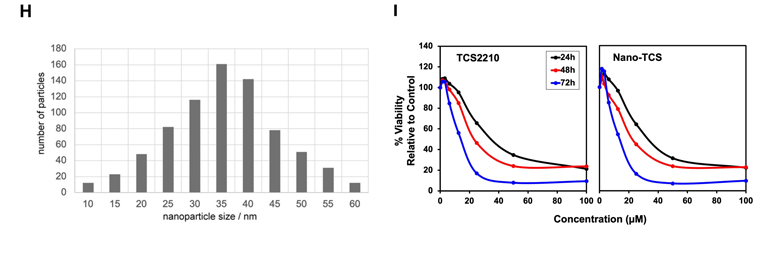
(A) Representative images of colony formation for A2780 and C30 cells treated with TCS2210. (B) Quantification of colony number of A2780 and C30 cells following treatment TCS2210. Data are presented as mean ± SEM from 3 independent experiments. (C) Quantification of colony size as in (B). Data are presented as mean ± SEM from 3 independent experiments. (D) Quantification of number of spheroids of A2780 and C30 cells treated with TCS2210. Data are presented as mean ± SEM, showing all data points from 3 independent experiments. (E) Hexosaminidase viability assay of A2780 cells treated with TCS2210, cyclodextrin (CD), or TCS2210:CD formulation. (F) Structure of nano-TCS2210 (nano-TCS). (G) Representative electron-micrograph of nano-TCS particles. (H) Quantification of Nano-TCS particle size. (I) Hexosaminidase viability assay of A1847 cells treated with TCS2210 or Nano-TCS.

